# The impact of low-frequency genetic variants on serum protein levels

**DOI:** 10.64898/2026.03.17.712047

**Authors:** Heida Bjarnadottir, Thorarinn Jonmundsson, Hulda K. Ingvarsdottir, Elisabet A. Frick, Nancy Finkel, Joseph J. Loureiro, Lenore J. Launer, Thor Aspelund, Yanhua Chen, Elizabeth K. Speliotes, Anthony P. Orth, Albert V. Smith, Valur Emilsson, Vilmundur Gudnason, Valborg Gudmundsdottir

## Abstract

The mapping of protein quantitative trait loci (pQTLs) can provide molecular links between genotype and phenotype. Most such studies focus on common variants, but the effects of low-frequency (LF) variants remain underexplored. Focusing on cis-pQTLs, we integrated serum measurements of 7,596 proteins with genomic data, including LF variants (minor allele frequency [MAF] 0.1-1%), in 5,291 Icelanders to identify independent cis-pQTLs for 2,166 SOMAmers. Incorporating LF variants increased the number of detected genetic signals per protein, demonstrating widespread allelic heterogeneity in cis-acting regulation of serum proteins. LF pQTLs were enriched for coding variants in the respective protein-encoding gene, but also among distal secondary signals, revealing additional regulatory layers not captured by common variants alone. Proteins affected by common variant cis-pQTLs were more often secreted and exhibited tissue-specific expression, whereas proteins exclusively affected by LF variants were primarily from more constrained and biologically essential pathways. Expanding both protein coverage and the allele-frequency spectrum reveals a more complex and heterogeneous cis-regulatory architecture of circulating proteins.

## Introduction

Genome-Wide Association Studies (GWAS) have successfully revealed thousands of genetic variations associated with disease risk, paving the way for personalized risk assessments and tailored therapies^1^. However, most disease-associated variants are in non-coding regions of the genome^2^, making functional interpretation difficult. Integration of high-throughput molecular data, such as transcriptomics or proteomics, can therefore provide insights into causal candidates mediating genetic effects on disease risk.

Studies have shown that incorporating human genetic evidence into drug development doubles the success rate from clinical development to approval^3^. Moreover, drug programs based on associations with rare genetic variants (MAF < 0.1%) or those exhibiting large effect sizes (odds ratio > 2) are three to four times more likely to achieve regulatory approval than those targeting common variants with small effects^4^. Despite challenges such as the potential for false positives and complex linkage disequilibrium (LD) patterns around rare variants, these findings underscore the importance of incorporating rare and low-frequency (LF) variant analysis into drug discovery and personalized therapies. With the recent development of high-throughput proteomics platforms, such as the aptamer-based SomaScan platform or the Olink immunoassay, it is now possible to simultaneously measure the circulating levels of thousands of proteins. Large-scale proteogenomic studies have identified thousands of protein quantitative trait loci (pQTLs)^5–13^, primarily driven by common variants, emphasizing the widespread genetic regulation of circulating protein levels. Such studies have driven the surge of protein Mendelian randomization (MR) analyses that leverage protein quantitative trait loci (pQTLs) to infer causal relationships between proteins and human diseases, identifying putatively causal protein-disease relationships and highlighting their value for drug target validation and prioritization^14^. Integrating MR with colocalization further refines these inferences by ensuring that protein and disease associations share the same underlying causal variant, thereby strengthening causal interpretation.

Historically, GWAS have primarily focused on common variants^15^. However, more recent reference panels, such as TOPMed^16^, have substantially improved imputation of rare (MAF < 0.5%) and LF variants compared to previous reference panels like 1000 Genomes^17^ and Haplotype Reference Consortium^18^. In admixed African and Hispanic/Latino populations, TOPMed increases the number of well-imputed rare variants by 2.3–6.1-fold and improves average imputation quality by 11–34%. Even extremely rare variants (MAF < 0.1%) show high information content recovery (>86%)^16,19,20^. This enhanced imputation enables the discovery of novel associations between LF variants and complex traits that would not be detectable with other panels.

While common variants are valuable for understanding the overall genetic architecture of circulating protein levels, LF variants can have a more substantial impact on protein function or regulation despite being less frequent in the population, as they often occur in critical regions of the genome that are less tolerant to change^21^. Studies of LF and rare variant effects on the circulating proteome have begun to emerge but remain scarce^11,22–24^. We have previously described numerous genetic and disease-associated patterns in the serum proteome from the Age, Gene/Environment Susceptibility (AGES)-Reykjavik Study cohort^8,25–28^, providing indications on causal chains from genetic variation to blood-protein levels to outcome. However, around third of the measured serum proteins have no identified common variant cis-pQTLs^8^, and we have shown that common variants tend to affect proteins located at the periphery in molecular networks, which may consequently have smaller individual effects on disease risk^8^. This observation is consistent with previously described systematic differences between genetic variants affecting disease susceptibility and gene expression, respectively, where disease-associated variants are biased towards more constrained genes and complex regulatory effects while eQTLs show the opposite pattern^29^. Thus, information on the genetic effects on central or “hub” proteins, the perturbation of which may be more important for disease susceptibility^30^, is lacking.

To address this issue, we here extend our previous GWAS of ∼5000 SOMAmer serum levels in the AGES cohort of elderly Icelanders^31^ with an updated SomaScan 7k platform (v4.1) by including 2,961 additional SOMAmers and a particular focus on LF (MAF of 0.1-1%) cis-acting variants. Cis-pQTLs are of particular interest as these may often reflect direct regulatory effects and offer greater interpretability than trans-acting associations. We identify cis-pQTLs for 693 SOMAmers not previously reported for similar cohorts^10,32^, thus markedly improving the coverage of proteins that can be tested in MR or colocalization studies. Importantly, we find 155 SOMAmers that are exclusively affected by independent LF cis-pQTLs and demonstrate that they are functionally more similar to proteins lacking any detectable genetic association than proteins influenced by common variants. LF variants also dominate distal secondary cis-signals, highlighting regulatory effects that remain hidden when only common variants are considered. Our results provide a catalog of genetic effects on functionally distinct sets of proteins that can be investigated for their phenotypic consequences and highlight the importance of including LF and rare variants in future proteogenomic studies. Full genome-wide association summary statistics from our study are made publicly available to facilitate their use in future proteogenomic studies.

## Results

### Proteome- and genome-wide association study

We measured the serum levels of 7,596 SOMAmers (Table S1) using the SomaScan 7k (v4.1) platform in 5,291 individuals with genetic data from the population-based AGES cohort (Fig. 1, Table S2). This analysis expands on our previous GWAS of 4,782 SOMAmers^8^, measured with a custom v3-5k SomaScan panel, of which 4,635 were also included in the 7k platform. The addition of 2,961 SOMAmers in the 7k SomaScan platform thus substantially broadened the scope of the previous study (Fig. 2A). The SomaScan assay exhibits exceptional sensitivity for the detection of low-abundance proteins, including intracellular proteins^33^. Consistent with this expanded coverage, the newly added SOMAmers were significantly enriched for proteins involved in RNA processing and biosynthesis (Supplementary Figure 1). Furthermore, tissue-specific expression patterns of the newly added SOMAmers pointed to enrichment in organs such as the duodenum, small intestine and kidney, highlighting potential roles in tissue-specific molecular functions and physiological regulation (Tables S3a-b).

**Figure 1:**
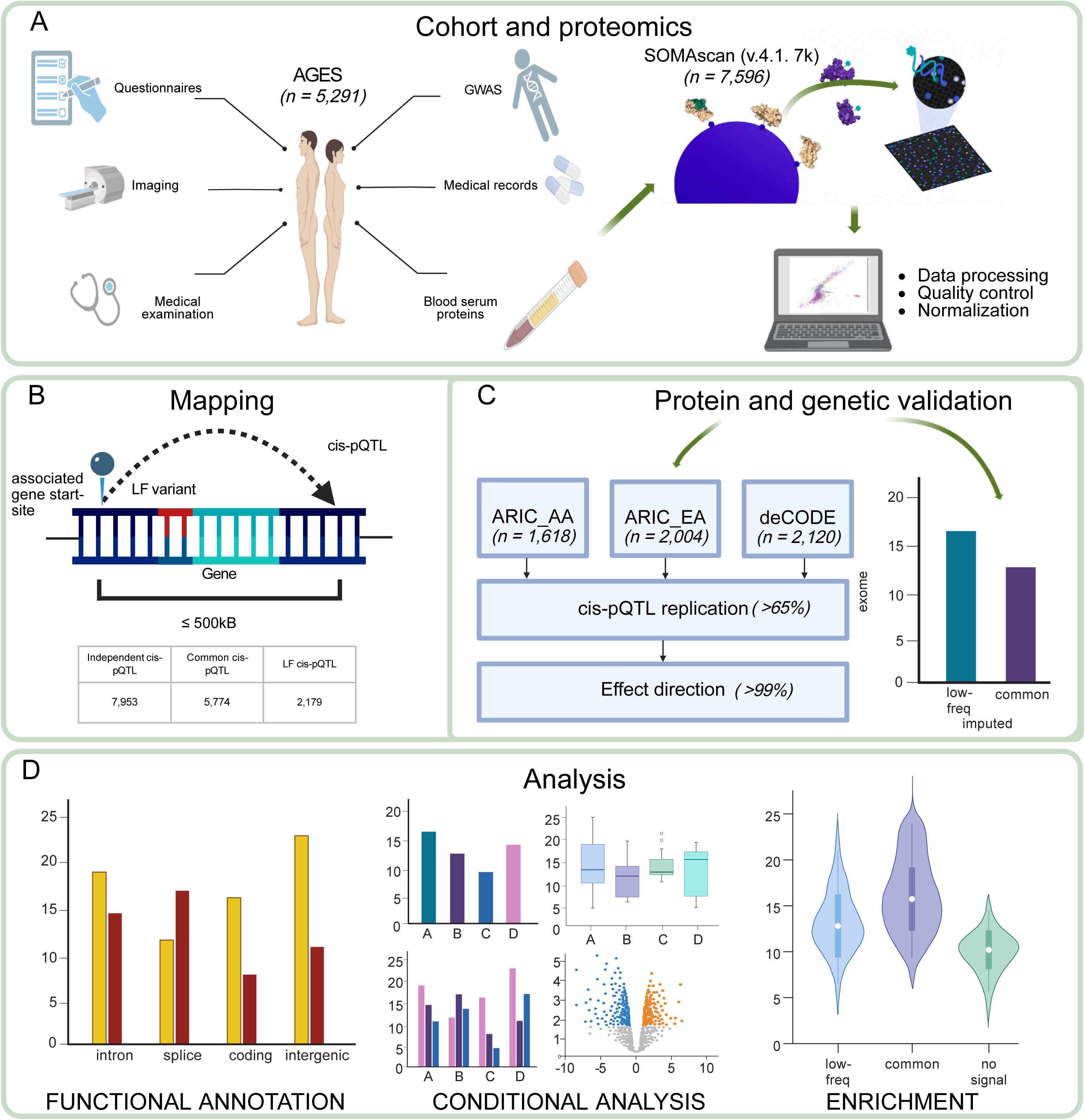
Study overview and analytical workflow. Low-frequency variants were defined as having minor allele frequency [MAF] 0.1-1%. **(A)** Deep proteomic profiling of the AGES cohort using SomaScan v4.1. (B) Cis-pQTL fmapping performed with QTLtools on TopMed-imputed genotypes (±500 kb window). (C) Protein-QTL validation in ARIC (AA/EA) and deCODE, including cis-replication and effect-direction. (D) Downstream analyses: functional annotation, stepwise conditional mapping, and enrichment of LF-only vs common signals.

**Figure 2:**
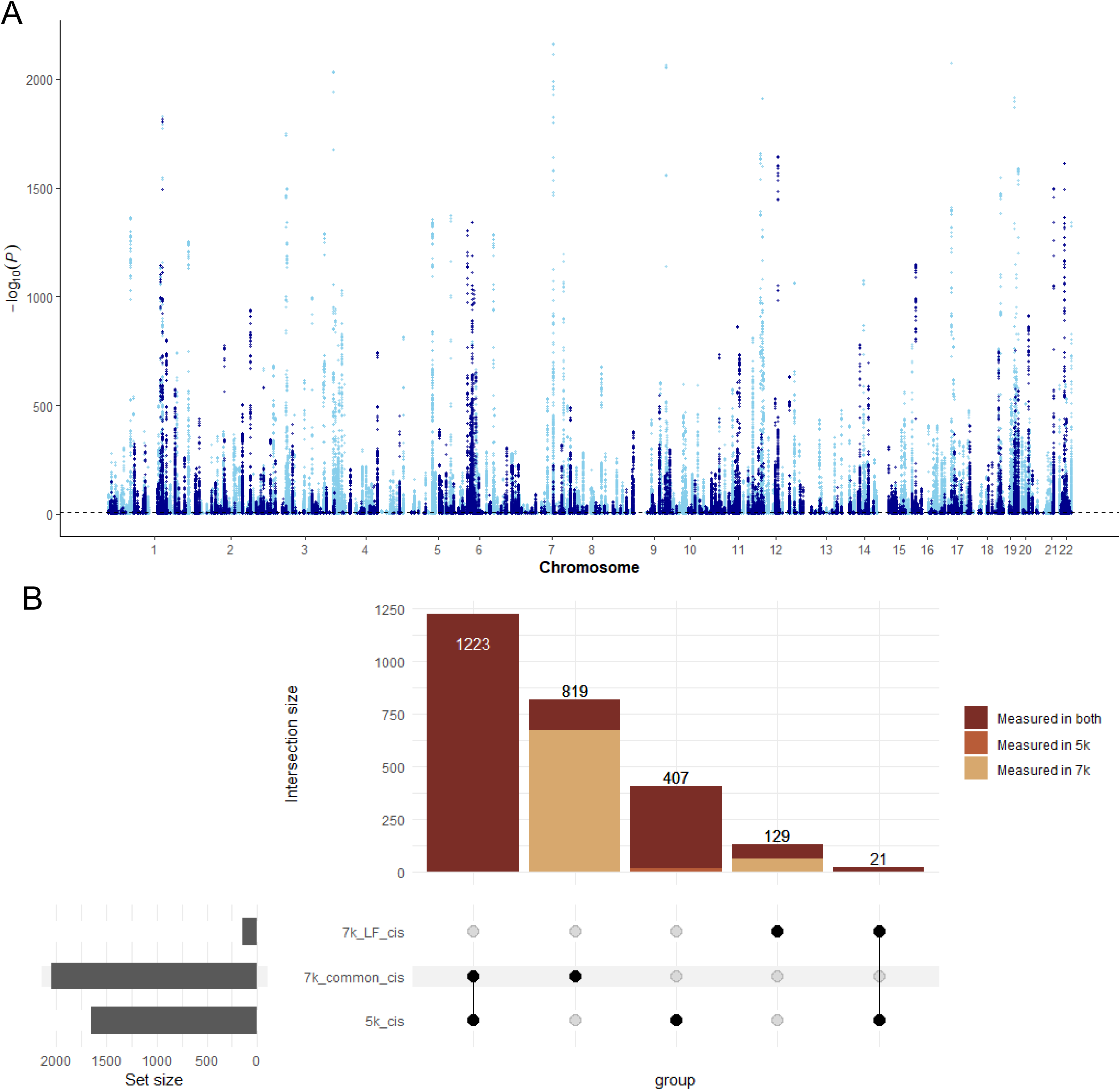
Overview of an expanded proteomic coverage and cis variant analysis using the 7k SomaScan platform. **(A)** A Manhattan plot representing the cis-associations detected using the 7k SomaScan platform. Points are colored by whether the corresponding protein target was already present in the earlier 5k platform (light blue) or newly included in the expanded 7k panel (dark blue). **(B)** UpSet plot showing the overlap of proteins with significant cis-pQTLs from the AGES 5k dataset and common or low-frequency cis-pQTLs from the AGES 7k dataset. Bars are colored by whether the protein was measured in both platforms, only in 5k, or only in 7k.

After quality control, the AGES genetic data included 11,919,579 variants (MAF > 0.1%) based on imputation to the TOPMed r2 reference panel, compared to the previously used 7,506,463 common variants (MAF > 1%) imputed to the Haplotype Reference Consortium (HRC) reference panel^8^. To validate the imputation quality of LF variants, we compared imputed genotypes for 47,747 variants that were also directly genotyped with the HumanExome BeadChip^34^ in 5,632 AGES participants. The genotyped variants showed consistent genotype calls when compared against the TOPMed-imputed data (Table S4), with LF variants averaging a concordance of approximately 0.999 (SD = 0.0017) over 11,876 sites, underscoring near-perfect agreement for these rarer alleles. Common variants, while exhibiting similarly strong concordance pattern, presented a slightly broader distribution of concordance values, averaging around 0.989 (SD = 0.0143) for 10,864 variants (Supplementary Figure. 2). These findings confirm that the genotyped and TOPMed-imputed genotypes are highly compatible, especially for LF variants.

We performed a GWAS on all 7,596 SOMAmers levels, identifying 1,245,212 variants for 4,072 unique SOMAmers with any study-wide significant associations (P < 5 × 10^-8^ / 7,596 SOMAmers = 6,582 × 10^-12^) (Fig. 2A). The average genomic inflation factor was low (mean λ = 1.035, SD = 0.031) implying minimal population stratification. Significant cis-pQTLs (defined as within 500 kb up- or downstream of the gene start and stop sites, respectively) were identified with a permutation-based empirical thresholding approach using QTLtools^35^, where the cis-significance threshold for each SOMAmer was determined based on 1,000 permutations (Methods). Sentinel variants reached the cis-pQTL significance criteria for 2,192 SOMAmers, of which 359 sentinel variants were LF while 1,833 were common (Table S5).

### Comparison of cis-pQTLs to previous studies

In our previous study, we identified 1,651 SOMAmers with window-wide significant common-variant (MAF > 0.01, P < 0.05/#SNPs per window) cis-pQTLs in AGES^28^, 1,244 of which overlapped with the 2,192 SOMAmers with cis-pQTLs identified in the current study (Fig. 2B). Thus, the 7k platform’s broader coverage enabled identification of 948 additional cis-signals, improving detection and allowing measurement of proteins not captured on the 5k platform (Fig. 2B). Of the 407 cis-SOMAmers previously identified in the 5k dataset but not replicated in the 7k analysis, 388 (95%) were measured on the 7k platform but did not meet the more stringent significance criteria, while 19 were not measured on the 7k platform. Among platform-shared SOMAmers, the proportion with significant cis-pQTLs was lower in the 7k dataset (29.5%) compared to the 5k dataset (35.2%), likely reflecting the stricter permutation-based significance threshold applied in this study. The average correlation of SOMAmer levels between platform versions was moderate (Pearsons, mean R^2^ = 0.66), potentially reflecting technical variation and updated assay properties between the v3 and v4 versions of the SomaScan platform, which could also contribute to the observed differences (Supplementary Figure 3A). Effect size estimates and p-values were highly correlated for sentinel SNPs between platforms (Spearman, ρ = 0.98 and ρ = 0.88, respectively), indicating good overall agreement (Supplementary Figures 3B–C). Finally, compared to our previous analysis of the HumanExome BeadChip (n variants = 54,469, MAF > 0.001) and the 5k SomaScan platform in AGES^13^, the current study adds information on cis-pQTLs for 1,394 additional SOMAmers.

We next compared the significant cis-pQTLs for the 7k platform in AGES with results from the deCODE (n = 35,892)^32^ and ARIC (African American [ARIC-AA]: n = 1,871, and European American [ARIC-EA]: n = 7,213)^10^ cohorts, both of which used the 5k (v4.0) SomaScan platform. This comparison revealed both shared associations, with 1,208 cis-SOMAmers shared between all four studies, and novel discoveries in AGES, or 693 cis-SOMAmers that were unique to the current study (Fig. 3A). The majority of these novel cis-SOMAmers (n = 590) were not measured in the comparison studies, due to the different version of SomaScan panel used, while 50 were identified only through the inclusion of LF variants.

**Figure 3:**
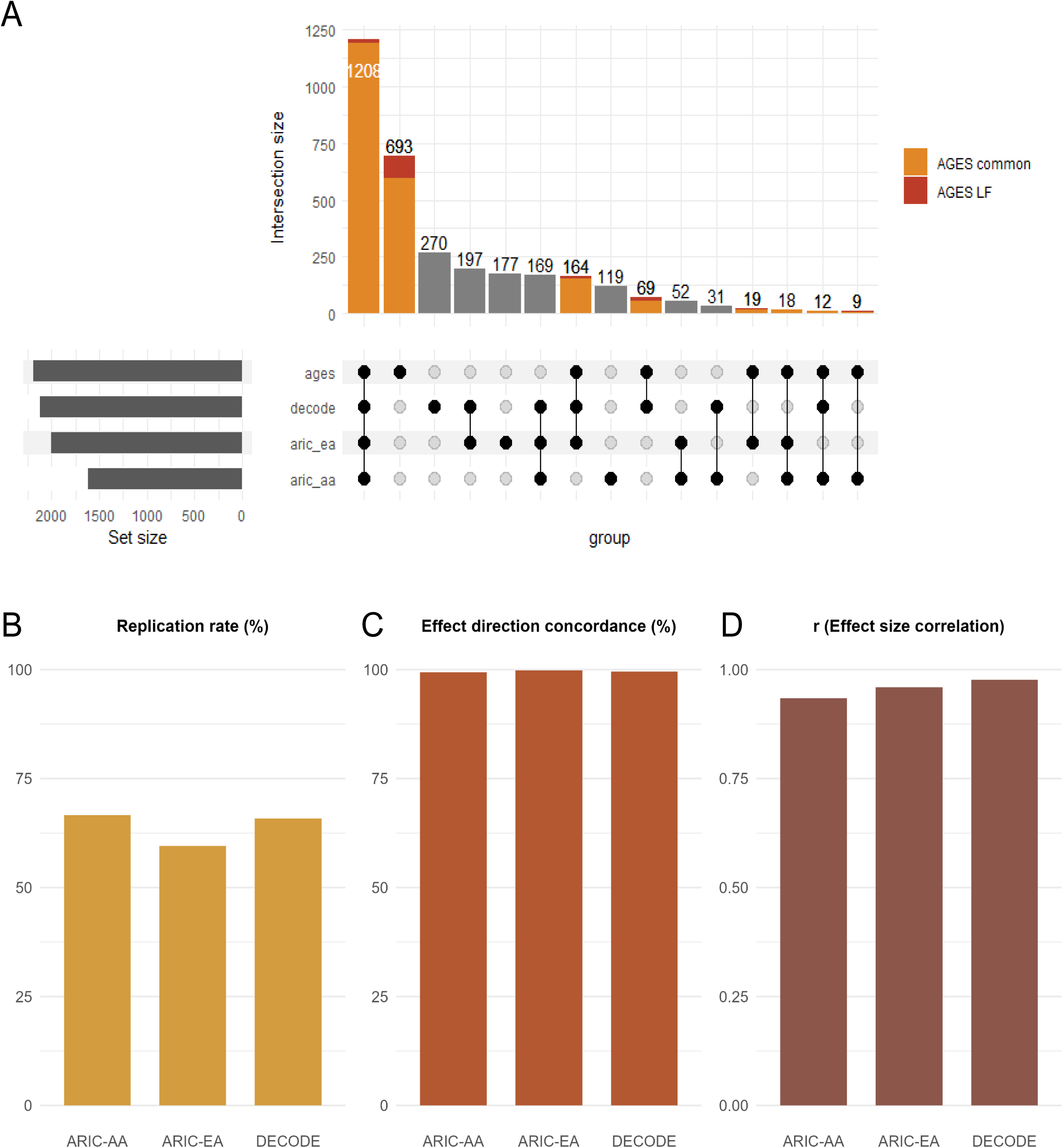
Cross-platform and cross-cohort replication of cis-pQTLs. **(A)** UpSet plot showing the overlap of cis-pQTLs in the AGES 7k dataset and the three external datasets: deCODE, ARIC-EA and ARIC-AA (SomaScan v4). Bars are colored by whether the signal was driven by common or LF variants in AGES. **(B)** Replication rate of the cis-pQTL from external datasets that reached a P < 2.36 × 10-5. **(C)** Effect direction concordance between replicated cis-pQTLs in AGES and the external datasets **(D)** Spearman’s correlation of effect sizes between AGES and external datasets for replicated cis-pQTLs.

Replication of reported cis-pQTLs from the deCODE and ARIC studies, which identified between 1,618 and 2,120 cis-pQTLs, was examined within AGES. The replication significance threshold was set using a Bonferroni correction based on the number of sentinel signals from the deCODE dataset (P < 0.05 / 2,120 = 2.4×10^-5^), which had the largest number of cis-pQTLs and therefore defined the most stringent multiple testing burden (Table S6). Of the 4,450 pQTLs that were replicable in AGES, replication rates of 65.7%, 59.3% and 66.6% were observed for the deCODE, ARIC-EA and ARIC-AA cis-pQTLs, respectively (Fig. 3B). Effect direction concordance was also evaluated for replicated signals, revealing > 99% concordance between AGES and all three external cohorts (Fig. 3C). We also found a strong correlation between absolute effect sizes in the AGES dataset and both the deCODE (r = 0.98) and ARIC-EA (r = 0.96) cohorts, while the correlation with the ARIC-AA cohort was slightly lower (r = 0.93) (Fig. 3D).

Of the 693 sentinel variants for the novel cis-pQTLs identified in AGES compared to the deCODE and ARIC studies, 514 (74%) had no reported proteomic associations (P < 1 × 10^-5^) in the GWAS-catalog^36^ (Table S7), supporting the novelty assignment of these pQTLs. The sentinel pQTL variant lookup additionally revealed 74 reported associations with non-proteomic traits, which in some cases implicated proteins other than those encoded by the nearest gene. Such examples include the *MTCH2* intron variant rs3817334, which has been consistently associated with body mass index (BMI) in multiple studies^37^. In AGES, this variant was associated with circulating levels of C1QTNF4 (C1q And TNF Related Protein 4), a recently discovered adipokine implicated in adipocyte biology, making it a biologically plausible candidate for mediating the obesogenic effects observed at this locus. C1QTNF4 is expressed in both brain and adipose tissue and has been shown in mouse models to directly influence food intake and energy balance^38^. The GWAS and pQTL evidence suggests this adipokine may also causally influence obesity in humans, highlighting it as an intriguing target for further investigation. Another example is the variant rs2896395, that is linked to insomnia^39^. While this variant is located in an intronic region of *SND1*, in AGES it was associated with serum levels of LRRC4 (Leucine Rich Repeat Containing 4), a synaptic adhesion protein primarily expressed in neurons. In mice it has been shown to play an important role for auditory synaptic responses mediating acoustic startle^40^ and thus has a physiological role that is in line with its observed GWAS association. Thus, the expanded pQTL mapping for circulating proteins presented here offers new opportunities to derive functional insights into disease-associated variants.

### Conditional analysis

To identify independent cis-pQTLs, we performed a conditional analysis using QTLtools^35^ (Methods). This resulted in 7,953 independent cis-signals (5,774 [72.6%] common and 2,179 [27.4%] LF sentinel variants, Fig. 4A) across 2,166 SOMAmers, corresponding to 1,893 unique proteins. The majority of the independent cis-signals, or 65%, was located outside the boundary of the respective protein-coding gene (Table S8).

**Figure 4:**
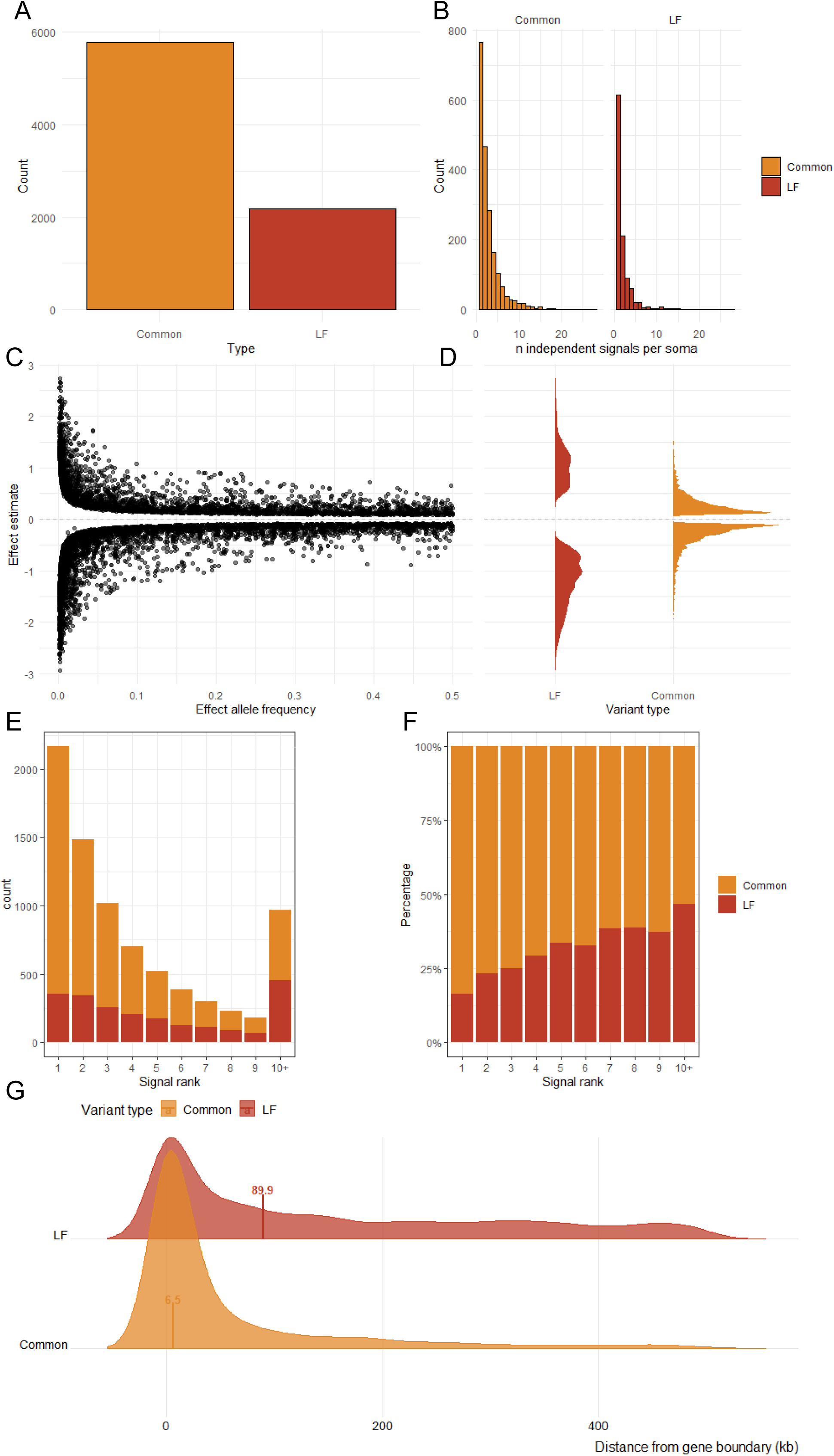
Summary of independent cis-pQTL signals from conditional analysis and explained variance. **(A)** Number of independent signals identified for common (MAF > 1%) and LF (MAF ≤ 1%) variants. **(B)** Distribution of the number of independent cis-signals per SOMAmer, stratified by variant type. **(C)** Distribution of effect size estimates of independent signals as a function of MAF **(D)** Distribution of effect sizes stratified by variant type, showing the negative skew and density of effect directions. **(E)** Counts of independent cis-pQTLs by signal rank, separated by variant type. The inset boxplot compares the distribution of signal ranks between common and LF variants, showing that LF variants tend to occur at later ranks. **(F)** Proportion of common versus LF variants within each signal rank. **(G)** Density distributions of independent cis-pQTLs by distance from the gene boundary, shown separately for common and LF variants. LF variants are typically located further from the gene, higher median distance (89.3 kb) compared with common variants (6.5 kb).

The number of independent signals per SOMAmer varied widely, from 1 to 46, with 1,482 SOMAmers (68%) carrying at least one secondary signal (Fig. 4B, Supplementary Figure 4A). LF variants contributed substantially to this allelic heterogeneity, as 49% of the cis-SOMAmers (1,053 of 2,166) harboured at least one independent LF signal. In fact, 155 SOMAmers were affected by LF-signals only, 35 of which were not captured in the initial permutation pass, reflecting that conditional analysis can uncover additional associations that are masked by stronger variants in the initial forward scan due to LD, multicollinearity or related effects. Among the 764 SOMAmers that appeared to have a single cis-pQTL when considering only common variants, 216 (28%) were reclassified as having multiple independent signals once LF variants were included (Supplementary Figure 4B).

At the variant level, independent LF variants showed a greater median effect absolute sizes (median |β| = 1.10, IQR 0.79-1.47) than common variants (median |β| = 0.24, IQR 0.15-0.41), which was also anticipated given the reduced statistical power for LF variants. Both common and LF variants had effect size distributions significantly skewed away from zero towards negative effects (Wilcoxon signed-rank test, Common: P = 1.01 × 10^-17^; LF: P = 1.78 × 10^-52^), with the LF variants having a stronger directional bias (Mann-Whitney U test, P = 1.31 × 10^-83^) (Fig. 4C-D).

Ranking the independent cis-signals per SOMAmer in the order of discovery, we observed that LF variants were increasingly represented among higher-ranked secondary signals, while primary signals were dominated by common variants (Fig. 4E-F). Primary signals had larger effects than secondary signals for both variant classes, but LF variants showed consistently larger absolute effects overall (Supplementary Figure 5). Higher-ranked signals tended to lie further from the gene boundary than primary signals with a gradual increase in median distance across rank groups (Kruskal-Wallis P = 1.25×10^-100^, Supplementary Figure 6). Consistently, LF signals also tended to be further away from the gene boundary than common variant signals (median ∼89 kb vs ∼5kb, respectively) (Fig. 4G).

Independent sentinel cis-pQTLs were distributed across multiple functional annotation categories, including coding, untranslated regions (5′ and 3′ UTR), promoters, and regulatory-region annotations (Fig. 5A). Enrichment was also observed for several transcript-processing categories such as splice-related annotations. The distribution of functional consequence categories differed significantly between LF and common variants (Chi-square[χ^2^] test, P = 6.0 × 10^-7^). Overall, LF variants were both more likely to be coding ([FET], P_adj_ = 2.4 × 10^-5^) and intergenic ([FET], P_adj_ = 0.04) than common signals, the latter reflecting the overall pattern of LF signals often capturing more distal secondary signals as described above (Fig. 5B). When restricting the analysis to cis-pQTLs located within gene boundaries of the respective protein-encoding gene, independent LF variants showed an even stronger enrichment for coding consequences ([FET], P_adj_ = 6.7 × 10^-39^) compared to common variants and were also enriched for splice-related annotations ([FET], P_adj_ = 2.3 × 10^-2^). In contrast, common intragenic variants were more likely to be non-coding, intronic, in UTR regions or Nonsense-Mediated Decay (NMD) transcript variants (Fig. 5C).

**Figure 5:**
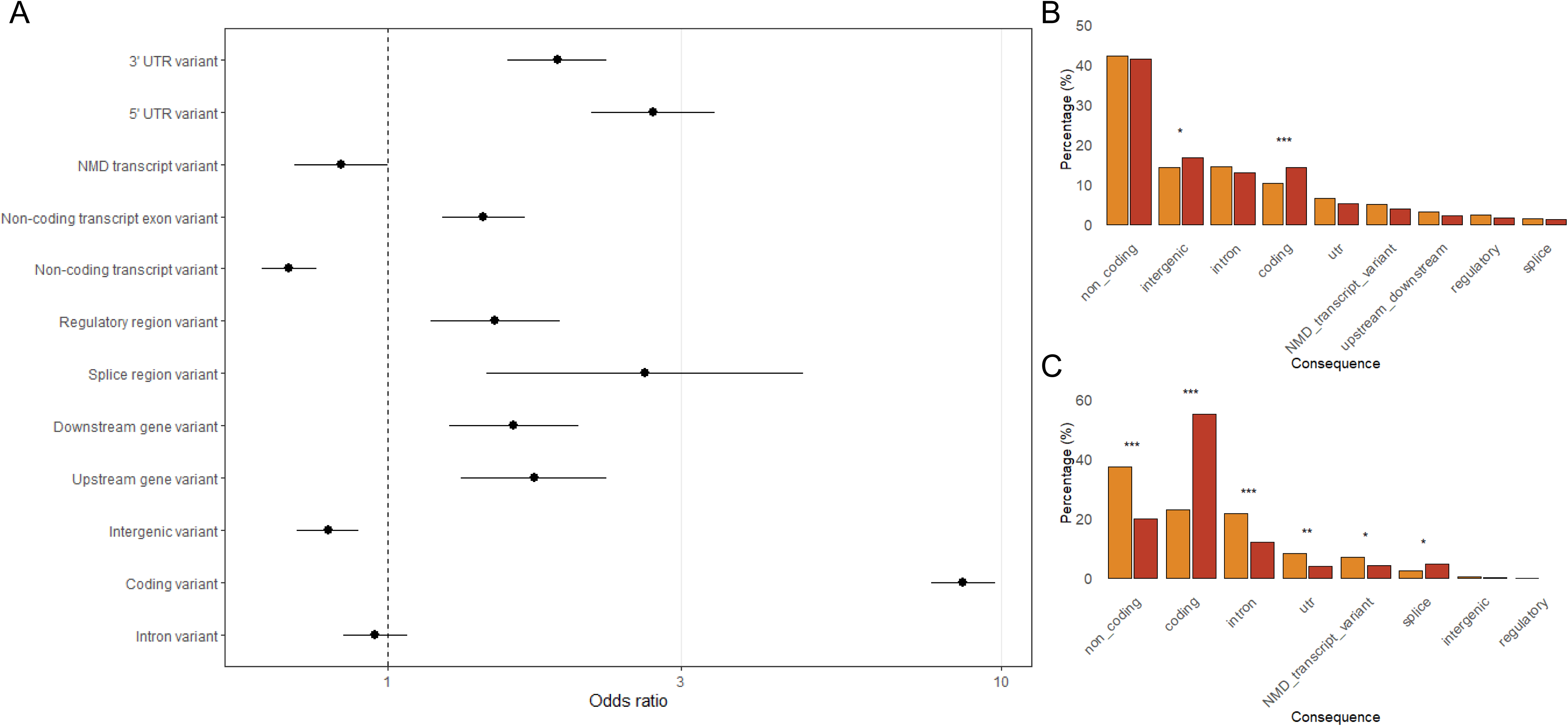
Functional annotation of independent cis-pQTLs. **(A)** Distribution of conditionally independent cis-pQTLs across Variant Effect Predictor (VEP v115) consequence categories. **(B)** Comparison of functional consequence categories between LF and common variants **(C)** Functional consequence distribution restricted to cis-pQTLs located within gene boundaries stratified by variant type (LF or common). **(A-C)** Significance was assessed with a one-sided Fisher’s exact test with Benjamini–Hochberg correction for multiple testing (*P < 0.05, **P < 0.01, ***P < 0.001; NS, not significant).

On average, common variants explained more variance in serum protein levels than LF variants. For 2,011 SOMAmers with at least one common variant cis-pQTL, common variants had a median R^2^*_Commo_*_n_ = 0.029 (IQR 0.009 – 0.1), compared to a median R^2^*_LF_* = 0.0095 (IQR 0.004 – 0.024) explained by LF variants for the 1,053 SOMAmers with any LF cis-pQTL (Fig. 6). However, for a substantial subset of SOMAmers, or 339 (15.7% of all SOMAmers with independent cis-pQTLs), LF variants contributed more to the explained variance than common variants (R^2^*_LF_* > R^2^*_Common_*) (Table S9). These included for instance CEL, SERPINA1, A1BG and APOA1, all of which are synthesized in the liver or intestines and play a role in metabolic and inflammatory pathways. Covariates contributed only modestly to serum protein level variance (Table S9).

**Figure 6:**
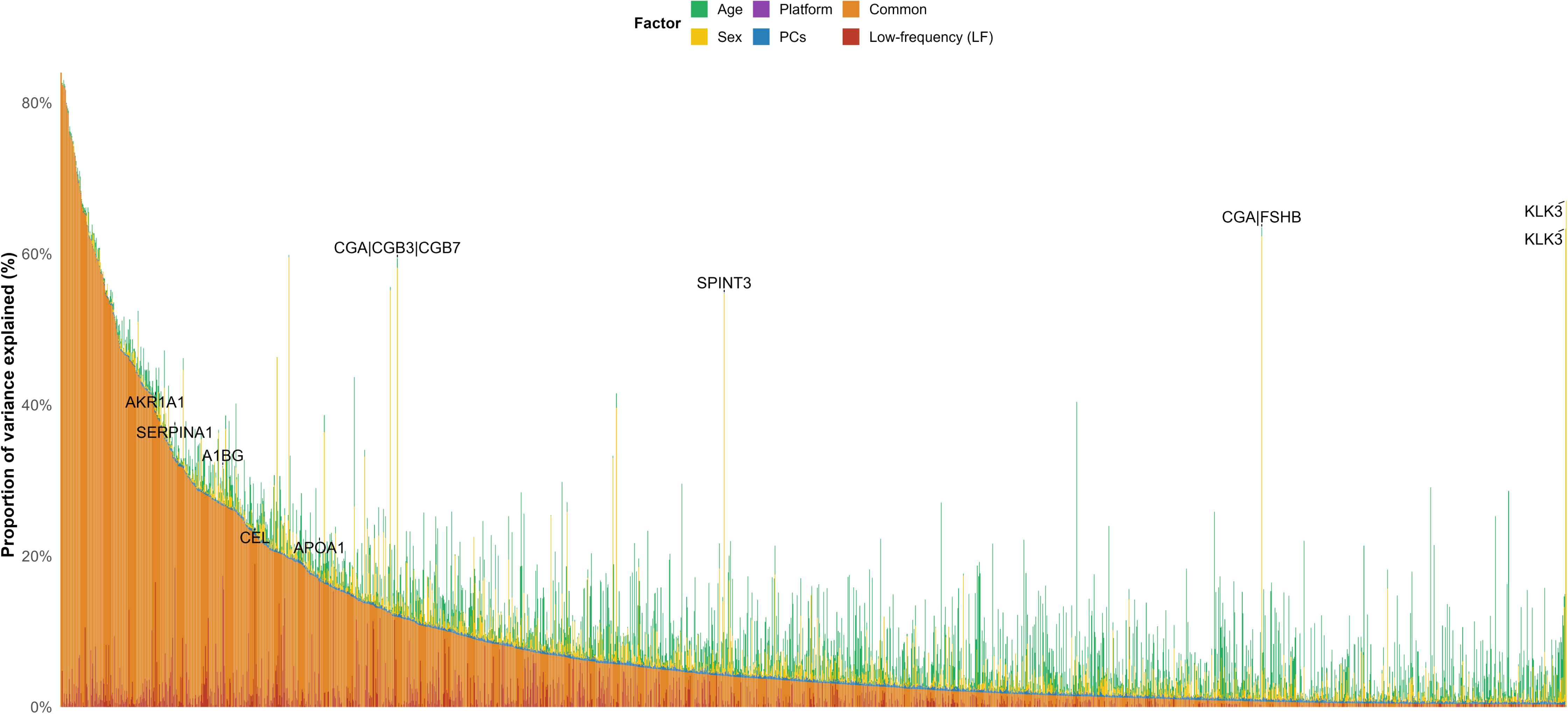
Proportion of variance in serum protein levels explained by conditionally independent cis-pQTLs and covariates. Proteins are ordered by decreasing variance attributable to genetics. Stacked bars show the relative contributions of LF and common genetic variants, as well as covariates (age, sex, genotyping platform and the first ten principal components).

### Characteristics of cis-proteins affected by LF-variants

To examine if the inclusion of LF variants would uncover cis-pQTLs for proteins with distinct characteristics compared to those identified in a common-variant analysis, an enrichment analysis was conducted at the level of unique proteins using conditionally independent cis-pQTLs. Proteins were classified into three mutually exclusive categories; a) any common cis-pQTL, b) LF-only cis-pQTL, and c) no cis-pQTL. Consistent with our previous study^8^, we found that proteins associated with any associated common cis-variants (n = 1,751) tended to have more tissue-specific gene expression profiles (FET, p_adj_ = 1.2 × 10^-20^) and were more likely to be secreted than other proteins (FET, p_adj_ = 3.2 × 10^-157^). In contrast, proteins exclusively affected by LF variants (n = 155) were less likely to have tissue-specific expression, exhibited a higher degree of connectivity in protein-protein interaction networks^41^ (Wilcoxon test, p_adj_ = 4.0 × 10^-5^) network^42^ and were less likely to tolerate loss-of-function (LoF) mutations (Wilcoxon test, p_adj_ = 4.0 × 10^-3^) (Fig. 7A-F). Overall, proteins associated exclusively with LF variants were functionally more similar to proteins lacking any cis-acting variant associations, indicating that including LF variants reveals genetic perturbations in proteins that would not be identified using common variants alone (Supplementary Figure 7).

**Figure 7:**
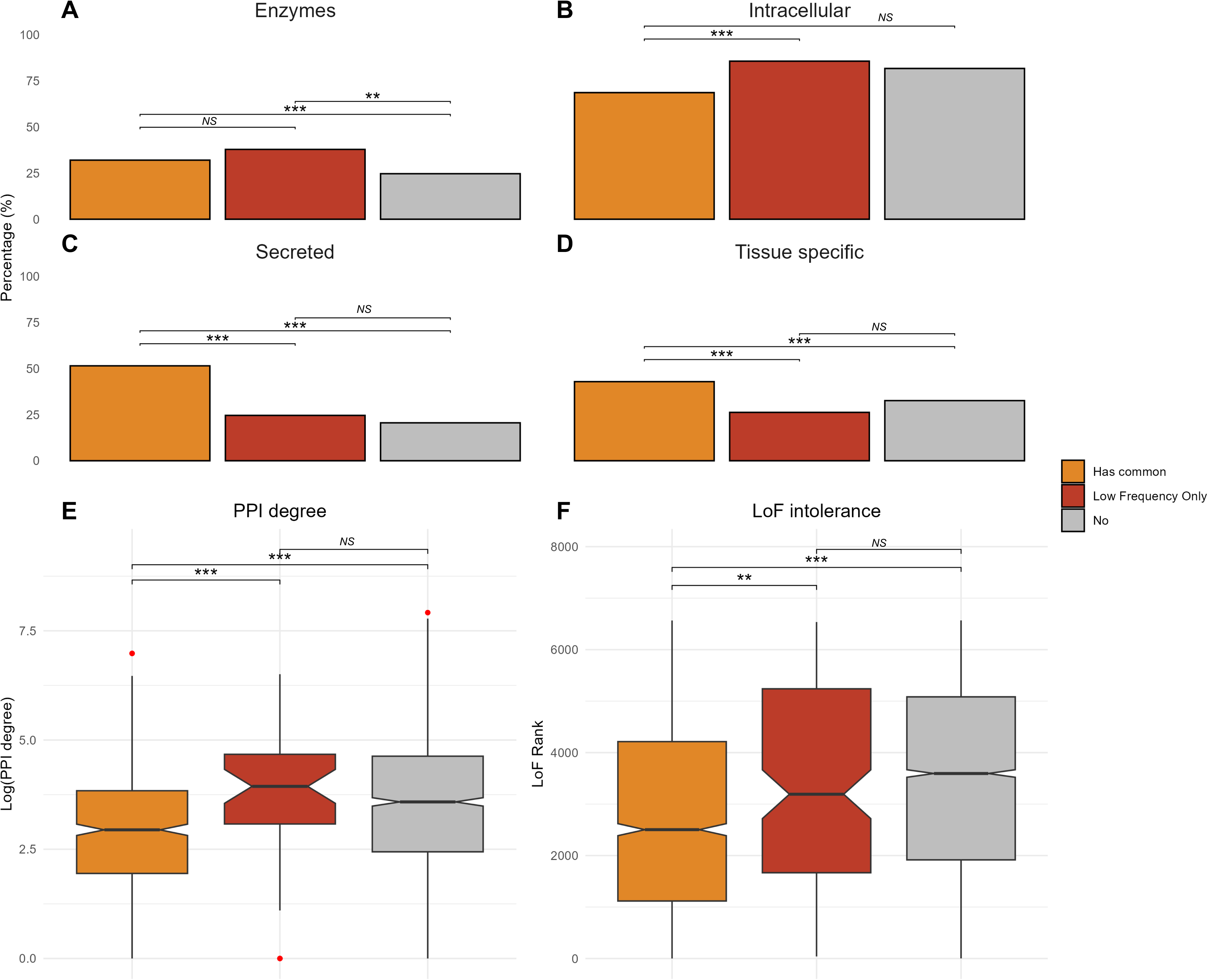
Protein class enrichment, PPI connectivity and LoF intolerance by cis-pQTL frequency category. Enrichment of **(A)** enzymes, **(B)** intracellular proteins, **(C)** secreted proteins and **(D)** tissue-specific gene expression among proteins with common cis-pQTLs (orange), LF cis-pQTLs only (red) or no conditionally independent cis-pQTLs (grey). Percentages indicate the fraction of proteins within each class and significance was assessed by pairwise Fisher’s exact test with Benjamini–Hochberg correction for multiple testing (*P < 0.05, **P < 0.01, ***P < 0.001; NS, not significant). **(E)** Degree of connectivity in the PPI network and **(F)** LoF intolerance rank for proteins with common, LF or no conditionally independent cis-pQTLs. **(A - F)** Significance was assessed by Wilcoxon rank-sum test (*P < 0.05, **P < 0.01, ***P < 0.001; NS, not significant). Boxplots indicate median value, 25th and 75th percentiles. Whiskers extend to smallest/largest value of no further than 1.5 interquartile range. Outliers are not shown.

Because we observed that LF variants contributed disproportionately to secondary cis-pQTLs, we investigated if these signals were generally detected for proteins with distinct biological properties. Among proteins with any cis-pQTL, having a secondary signal was associated with higher odds of the protein being secreted (FDR = 1.6 × 10^-19^) and having more tissue-specific expression (FDR = 2.03 × 10^-4^). After incorporating LF variants into the signal classification, these proteins also showed reduced tolerance to LoF variation (FDR P = 2.35 × 10^-3^) and were enriched for disease-associated genes (FDR = 8.41 × 10^-4^) (Fig. 8). Thus, while greater allelic heterogeneity was associated with less selective constraints, it was also indicative of disease relevance, in particular when incorporating information on LF variants.

**Figure 8:**
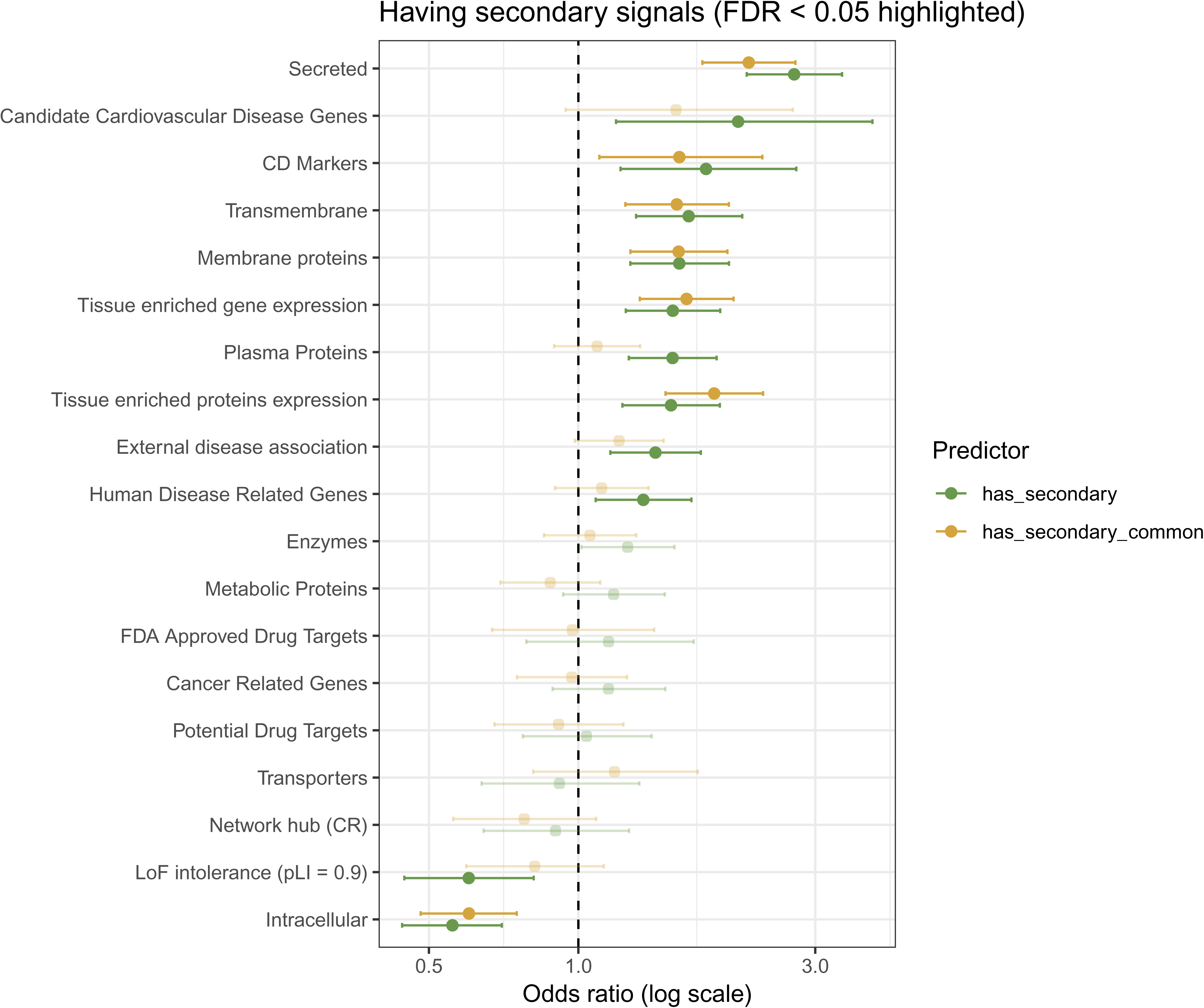
Enrichment annotations associated with the presence of secondary cis-pQTL signals. Forest plot showing odds ratios (log scale) from logistic regression models testing whether proteins with secondary cis-pQTLs (green) or secondary signals identified from common variants only (gold) are enriched for specific gene or protein annotations. Points represent estimated odds ratios and horizontal bars show 95% confidence intervals and annotations having FDR < 0.05 are highlighted.

## Discussion

In this study, we systematically characterized the contribution of LF and common genetic variants to serum protein variation using conditionally independent cis-pQTL mapping. Incorporating LF variants substantially broadened the view of the genetic architecture of the circulating proteome and increased the number of independent signals per protein, particularly distal secondary signals, highlighting widespread allelic heterogeneity in cis-acting regulation. Importantly, including LF variants revealed associations for a distinct subset of proteins that would be undetected if the analyses were limited to common variants. We expanded the current cis-pQTL landscape by identifying 693 novel proteins with significant genetic associations that were not reported in either of the two external proteogenomic studies used for comparison^10,32^. This expanded catalog provides new opportunities for protein MR and colocalization analyses, enabling causal evaluation of newly identified proteins and systematic prioritization of potential therapeutic targets.

Extensive allelic heterogeneity was evident at the protein level. More than two-thirds of proteins carry at least one secondary cis-pQTL, and LF variants affect over half of proteins with multiple independent signals. We found that proteins with secondary cis-pQTL signals were enriched among disease-associated genes but were depleted among LoF-intolerant genes, consistent with the idea that highly constrained genes tolerate fewer independent regulatory perturbations^29^. While intragenic LF-pQTLs tended to be driven by coding variants, LF cis-pQTLs outside of the protein-encoding gene were located further away than common variants, similar to secondary signals overall. However, functional enrichment analyses did not provide a conclusive explanation for this distance pattern, as no differences were observed between common and LF in regulatory regions. However, a similar trend toward more distal localization of secondary signals with lower frequency has recently been reported in adipose eQTL analyses^43^, suggesting that this may reflect a general regulatory architecture. Importantly, the inclusion of more distal non-primary eQTLs was found to considerably improve colocalization with GWAS loci^43^.

Extending this paradigm to the proteome demonstrates that conditional mapping combined with an expanded allele-frequency spectrum can substantially increase the discovery of cis-pQTLs, thereby refining the landscape of genetically regulated proteins and broadening the set of potential molecular instruments for downstream applications, such as Mendelian randomization and drug target prioritization. An important next step will be to extend these analyses to trans-pQTLs, which may hold particular importance for GWAS loci^44^, where the effects of LF variants remain largely unexplored and could uncover additional long-range regulatory effects on protein networks that are not detectable through common-variant or cis-focused approaches.

We observed that proteins influenced primarily by common cis-pQTLs were more likely to be secreted and tissue-enriched, distinguishing them from broadly expressed housekeeping proteins and making them experimentally accessible and potentially useful in longitudinal monitoring studies. In contrast, proteins regulated by LF variants showed stronger connectivity within PPI networks, potentially pointing to functional complexes carrying out a specific biological role. These proteins also displayed less tolerance to LoF variants, consistent with being under stronger evolutionary constraint^45^. This contrast is consistent with the possibility that common variant associations primarily involve proteins whose expression can vary without compromising cellular function whereas LF cis-pQTLs affect proteins embedded in essential, tightly regulated pathways, where mutations carry a greater risk of functional disruption. Notably, proteins with LF-only signals resembled proteins without detectable cis-pQTLs in several characteristics, suggesting that rarer regulatory variation may act on more tightly controlled and less expression-flexible genes.

Replication analyses demonstrated overall strong consistency across cohorts, confirming the robustness of the cis-pQTL effects identified in AGES. However, several limitations should be considered. Minor differences in replication rates across AGES and external datasets likely reflect differences in sample sizes, population-specific linkage disequilibrium structures and allele frequency differences, while platform and analytical differences may also contribute modest variability. Furthermore, the slightly reduced correlation in effect sizes observed in ARIC-AA likely reflects the distinct genetic architecture of African ancestry populations^45^. Additionally, power to detect LF variants is limited, both statistically and by imputation accuracy, which could bias effect estimates or obscure true signals. Finally, while conditional analysis enables discovery of multiple independent cis-pQTLs, it relies on linear modelling and may not capture more complex regulatory architectures.

In conclusion, expanding both protein coverage and the allele-frequency spectrum reveals a more complex and heterogeneous cis-regulatory architecture of serum proteins than previously appreciated. LF variants contribute substantially to secondary signals and overall allelic heterogeneity, extend associations to more distal associations, and uncover regulation of proteins that would otherwise remain undetected. Our results provide an expanded framework for understanding genetic regulation of the circulating proteome and establish a foundation for downstream causal inference and target prioritization analyses.

## Materials and Methods

### AGES-Reykjavik study

The AGES-Reykjavik study is a single-center, population-based cohort comprised of 5,764 participants aged 66 - 96 years (mean age 76.6 ± 5.6 years; 58% women)^31^. This study builds on the long-term Reykjavik Study, aiming to understand the aging process through comprehensive assessments of four primary biological systems: vascular, neurocognitive (including sensory), musculoskeletal, and body composition/metabolism^46–49^. participants underwent extensive phenotypic evaluations, including anthropometric measurements, questionnaires and detailed clinical assessments as has been described previously^6,8,9,25–28,50^. Ethical approval for the study was obtained from the National Bioethics Committee in Iceland (VSN-00-063), the National Institute on Aging Intramural Institutional Review Board, and the Data Protection Authority in Iceland. All participants provided informed consent prior to enrolment.

### Genotyping and imputation

Within the AGES cohort, 3,219 individuals were genotyped using the Illumina hu370CNV array, and 2,710 individuals were genotyped using the Illumina Infinium Global Screening Array (GSA). To avoid duplicate data, 268 individuals overlapping between the arrays were excluded from the hu370CNV. Both datasets were imputed to the TOPMed reference panel, containing over 400 million variants from diverse populations^16^, via the Michigan imputation Server^51^. The genotypic data was initially pre-phased using Eagle v2.4 for accurate haplotype phasing. Imputation was performed with Minimac4. Genotypes from the hu370CNV and GSA platforms were merged based on matching locations and alleles. Quality control measures were applied post-imputation (and post merging) to ensure data reliability. Variants were excluded if they had MAF < 0.001, deviated significantly from Hardy-Weinberg equilibrium (HWE P < 1 × 10^-6^), or had an imputation quality score (R^2^) < 0.7. Multi-allelic variants, copy number variants (CNVs), and other non-SNP variants were excluded during this process which was conducted using BCFtools (v1.9)^52^ and PLINK 1.9^53^. All genomic positions were aligned to the GRCh38 reference genome.

To validate concordance between directly genotyped variants from the HumanExome BeadChip^34^ and imputed TOPMed data, overlapping SNPs were first extracted by matching on rsIDs (excluding multiallelic sites and duplicates). Genotypes from both datasets were merged, and per-variant concordance rates (allowing strand flips) were calculated as the fraction of non-missing calls that agreed. Concordance was then stratified into common (MAF > 0.01) and LF (MAF ≤ 0.01) bins, with mean and SD reported.

### Protein GWAS and detection of significant cis-SOMAmers

Within the AGES cohort, 5,291 individuals had both genetic data and serum protein level measurements at baseline. Using that sample set, 11,919,579 (of which 3,850,407 had MAF ≤ 0.1 - 1%) variants were tested for association with each of the 7,596 SOMAmers separately, using age, sex, 10 genetic principal components, and genotyping platform as covariates. A genome-wide analysis was performed using PLINK 2.0^54^, fitting linear models to Box-Cox transformed, centered and scaled protein levels. Extreme outlier values were excluded (defined as values above the 99.5th percentile of the distribution of 99th percentile cutoffs across all proteins after scaling, resulting in the removal of an average 11 samples per SOMAmer), as previously described^25^.

We performed cis-pQTL discovery using QTLtools (v1.3.1) with a ±500 kb cis-window, using the same covariates as in the genome-wide analysis. First, a nominal pass (--nominal 0.001) was run to obtain association statistics for all variants within the cis-window. Next, we applied a permutation pass with 1,000 permutations (–permute 1000) to estimate empirical significance while accounting for the LD structure among nearby variants. For each SOMAmer, permutations were used to derive the null distribution of the minimum association p-value across all variants tested in the cis-region. Permutation-derived p-values were adjusted using the beta approximation implemented in QTLtools, and the resulting beta-adjusted p-values were corrected across SOMAmers using the Benjamini–Hochberg false discovery rate procedure. For each SOMAmer, we classified whether the detected cis-pQTL signals were driven exclusively by LF variants, exclusively by common variants, or both.

### Comparison between platform versions and previous proteogenomic studies

To compare the cis-pQTLs identified across platform versions in AGES, we contrasted results from the earlier v3-5k GWAS^8^ (HRC imputation) with those from the current v4.1-7k platform (TOPMed imputation). The datasets were merged based on shared genomic positions and SOMA identifiers to identify overlapping and unique cis-pQTLs between the platforms. Effect sizes and p-values from the 5k and 7k datasets were compared and Spearman’s correlation coefficients were calculated to evaluate consistency between platforms. The number of unique cis-pQTLs detected due to the inclusion of LF variants on the 7k platform was quantified.

All cis-pQTLs identified on the 7k platform were compared with previously published cis-pQTLs from the deCODE^32^ and ARIC^10^ cohorts, the latter of which includes both African American (ARIC-AA) and European American (ARIC-EA) populations. Overlapping variants were matched across datasets by rsIDs and alleles, excluding multiallelic SNPs and duplicates. In the ARIC study, the cis windows was defined within ±500 kb of the associated gene start site while the deCODE study defined cis-pQTLs within 1 Mb of the transcription start site. For each cis-protein reported in the external datasets, the sentinel SNP was selected based on the lowest p-value within the cis-window. Effect sizes and p-values for the sentinel SNPs were compared across datasets. Replication of external cis-pQTLs in AGES was assessed using a significance threshold of P < 2.36 × 10^-5^, corresponding to a Bonferroni correction for 2,120 tests (the maximum number of cis-pQTLs identified in the deCODE dataset).

### Identification and classification of primary- and secondary cis-pQTL signals

Conditionally independent cis-pQTL signals were identified using the QTLtools^35^ (v1.3.1) conditional pass (--mapping) using a ±500 kb cis-window. QTLtools implements a forward-backward stepwise regression procedure, which accounts for local LD to detect multiple independent signals per SOMAmer. For each SOMAmer, all variants identified as backward-significant were retained, and the most significant variant (lowest p-value) was selected as the sentinel (primary) for each signal rank (rank 1). Independent signals were considered significant if their permutation-derived empirical p-value (from the –permute 1000 scan) passed a 5% FDR threshold. Proteins with more than one independent signal were classified as having secondary signals, and those with exactly one signal were classified as primary-only.

### Variance explained

Per-protein variance explained was quantified using block partial R^2^ from linear models fit to aligned samples. For each protein, the outcome was the protein level and predictors were grouped into blocks: (i) LF genotypes (all independent cis variants with MAF ≤ 1%), (ii) Common genotypes (MAF > 1%), and (iii) the covariates age, sex, genotyping platform, and the first ten genetic principal components.

### Variant annotation and functional enrichment

Variant annotation was based on Ensembl Variant Effect Predictor (v.115)^55^, classifying variants as non-coding, coding, intronic, intergenic, splice site, up- or downstream, untranslated regions (UTRs), or regulatory. For each variant, the most severe predicted consequence was retained. To assess functional enrichment of sentinel independent cis-pQTLs relative to non-pQTL cis-variants, we performed an LD-aware enrichment analysis using logistic regression. All cis variants were first LD-pruned (r^2^ < 0.1) to generate a set of seed variants. For each seed variant, LD proxies were identified within ±500kb at r^2^ > 0.8 using PLINK (v1.9). Functional annotations were assigned at the proxy level and aggregated to the seed level, such that a seed variant was considered annotated for a given feature if any of its proxies carried that annotation. For each feature, we modelled the probability of annotation (binary outcome) as a function of pQTL status (sentinel cis-pQTL vs. non-pQTL cis variant) using a logistic regression. Models were adjusted for allele frequency by including an indicator for LF status. To compare functional consequence profiles between LF and common sentinel cis-pQTLs, we compared the distribution of VEP consequence categories using two-sided Fisher’s exact tests, followed by Benjamini–Hochberg correction for multiple testing across tested categories. As a complementary global test, differences in the full contingency table of consequence categories by variant class were assessed using a χ^2^ test. To evaluate whether observed differences were driven by variant-to-gene localization, we repeated the LF-versus-common comparisons after restricting to cis-pQTLs located within gene boundaries.

### Protein level enrichment analysis

All measured proteins were categorized into three groups based on conditionally independent pQTLs; a) proteins with only LF cis-pQTLs, b) proteins with at least one common cis-pQTL, and c) proteins without any cis-pQTLs. For the enrichment analysis, these protein categories were compared against external features. These included annotations for protein characteristics from the Human Protein Atlas (HPA)^56^, gene^56^ and protein^57^ expression across tissues, LoF intolerance^58^, connectivity metrics from the InWeb protein-protein interaction network^42^ and the serum protein co-regulation network derived from the AGES cohort^6^. Disease association information was integrated from curated databases, including OMIM^59^, Monarch Initiative^60^ and Orphadata^61^, which provide established gene-disease (rare and common) and phenotype associations. One-sided Fisher’s exact test was employed to evaluate associations between protein categories and binary classifications. For continuous variables, the Wilcoxon rank-sum test was used to compare across protein groups. To account for multiple comparisons, the Benjamini-Hochberg procedure was applied to adjust the p-values considering FDR < 5% as statistically significant. Pathway enrichment analysis was conducted using gProfiler^62^, with the complete set of measured proteins used as the background. Statistical significance was determined based on a Benjamini–Hochberg adjusted threshold of < 0.05.

## Resource availability

### Data and code availability

- All original code has been deposited at Zenodo and is publicly available as of the date of publication. All statistical analyses were performed using R version 4.4.1 and Rstudio version 2025.05.0.
- All independent cis-pQTLs are provided in supplementary data, while full summary statistics are available for download.
- Any additional information required to reanalyze the data reported in this paper is available from the lead contact upon request.

### Competing interests

N.F. and J.J.L. are employees and stockholders of Novartis. The Regents of the University of Michigan and EKS have a pending patent on the use of systems and methods for analysis of samples associated with MASLD, insulin resistance, and related conditions.

## Acknowledgements

National Institute on Aging contracts N01-AG-12100 and HHSN271201200022C, the Icelandic Heart Association, and Althingi (the Icelandic Parliament) financed the AGES-Reykjavik study. IHA and Novartis have collaborated on proteomics research since 2012. The study was also funded by the Icelandic Centre for Research (grant no. 2511008-051) and the University of Iceland Research Fund (grant no. 94151). Research reported in this publication was supported by the National Institute On Aging of the National Institutes of Health under Award Number R01AG065596. The content is solely the responsibility of the authors and does not necessarily represent the official views of the National Institutes of Health. YC and EKS are supported by NIH grants R01 DK106621, R01 DK107904, R01 DK131787 (to EKS) and R01 DK128871 (to EKS), the University of Michigan Department of Internal Medicine and MBIOFar.

