## Supplementary Material for "The impact of low-frequency genetic variants on serum protein levels"

### **Supplementary Figures**

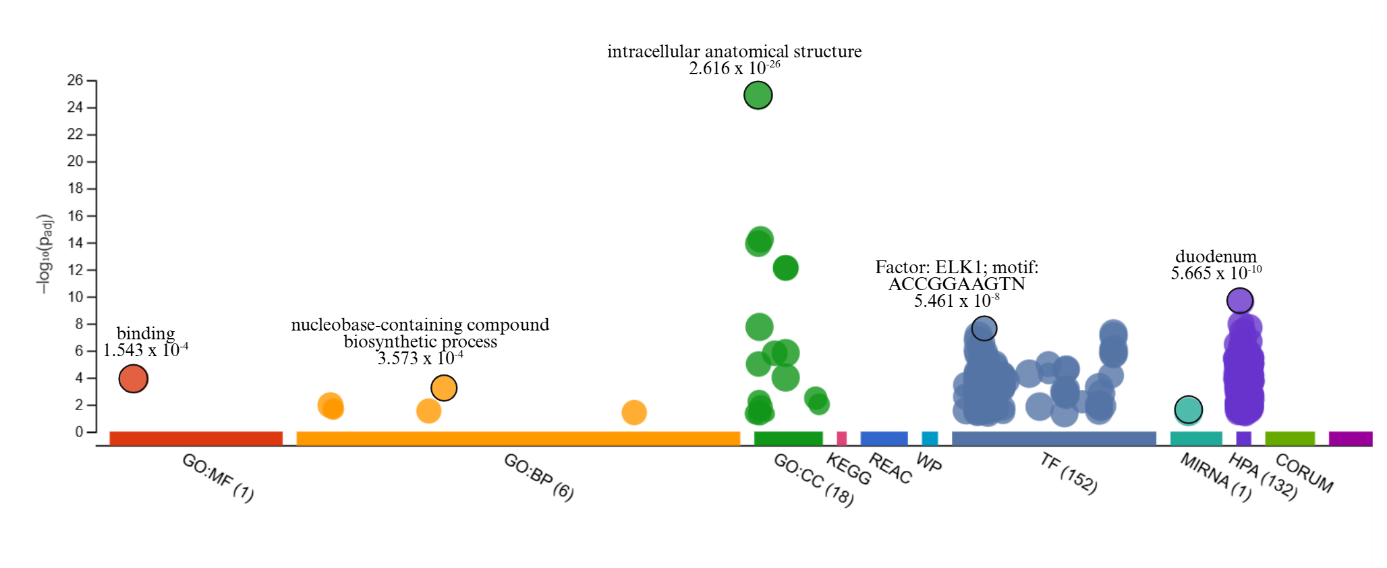

#### **Supplementary Figure 1 | Functional characterization of the 2,961 proteins uniquely included in the 7k SOMAscan platform.** Shown are enriched biological processes and molecular functions based on gene ontology annotations, KEGG, reactome and wikipathways.

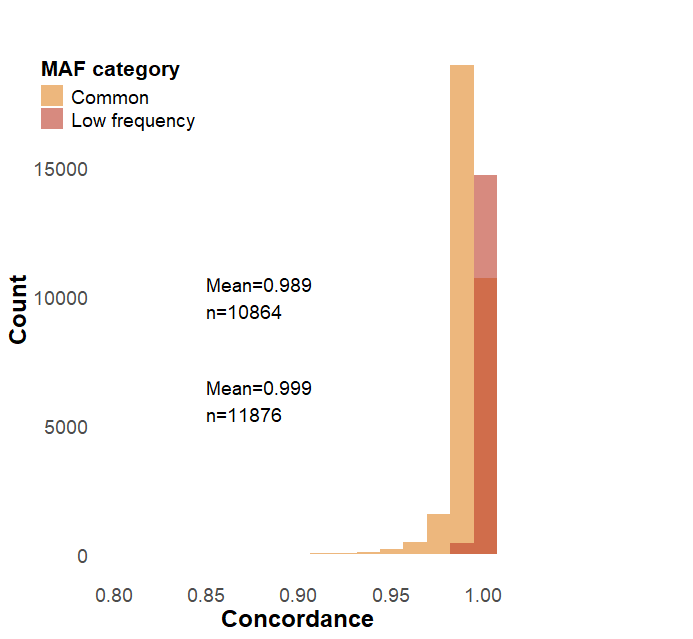

**Supplementary Figure 2 | Concordance of TOPMed-Imputed and Exome Chip Genotypes**. Both LF and common variant sets are tightly clustered near the upper limit of concordance, reflecting minimal discrepancies. Nevertheless, the wider spread in the common variant subset suggests that while concordance is still very high, there is a small fraction of variants with slightly reduced agreement between the two datasets

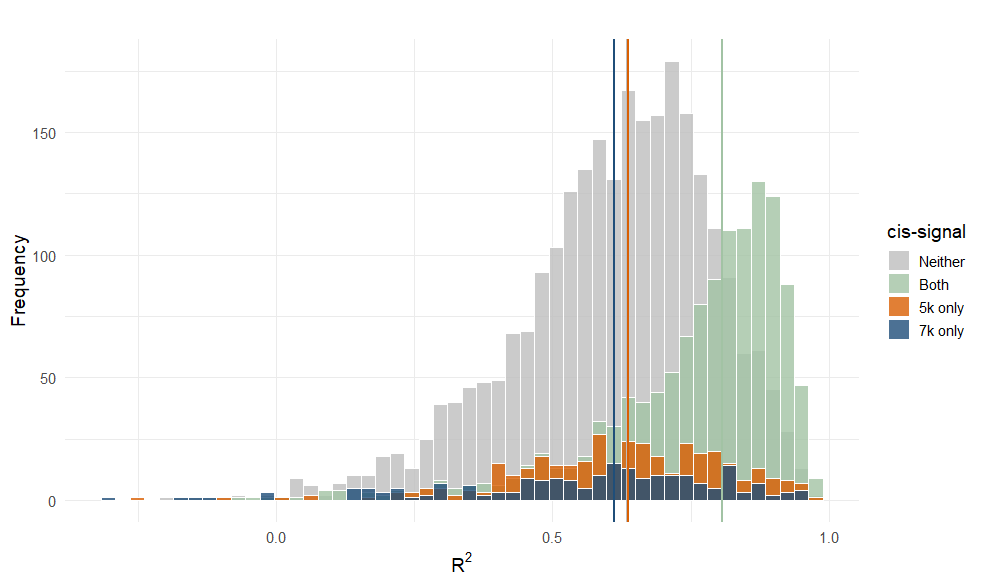

**Supplementary Figure 3.A | Cross platform correlation of somamer levels.** Distribution of squared Pearson correlations (R^2^) between protein measurements on the 5k (v3-Novartis) and 7k (v4.1) platforms for shared proteins, stratified by cis-pQTL status (“Both”, “5k only”, “7k only”, “Neither”). The average correlation was moderate (R^2^ = 0.65). Proteins with cis-pQTLs detected in both platforms show higher measurement agreement than proteins with platform-specific or no cis-pQTLs (median: R^2^ neither: 0.634, both: 0.806, 5k only: 0.637, 7k only: 0.612).

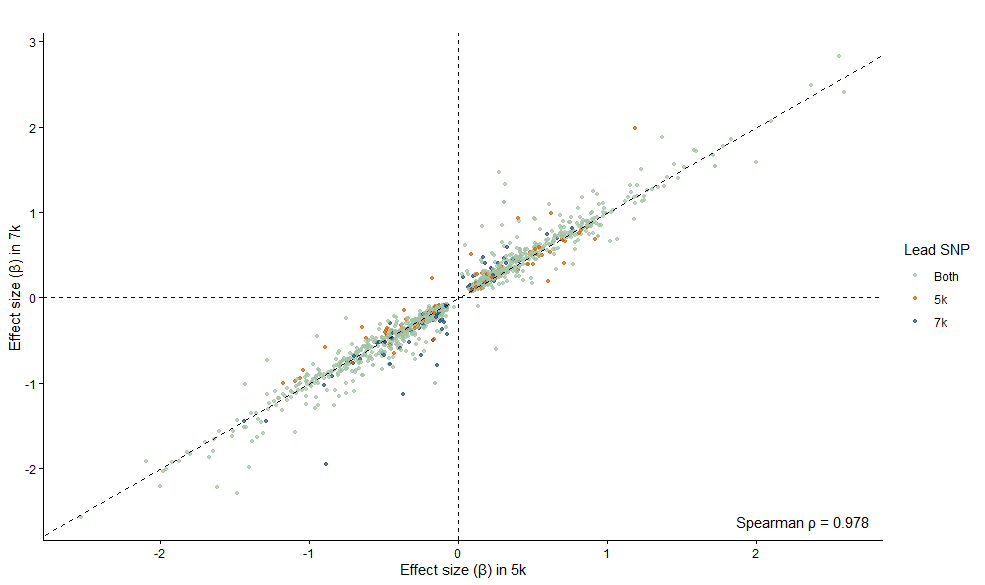

**Supplementary Figure 3.B | Correlation of cis-pQTL effect sizes between 5k (v3-Novartis) and 7k (v4.1) SOMAscan platforms.** Each point represents a variant-protein pair tested within the cis window and present in both datasets. Points are colored by whether or not the signal is detected in 7k (green), 5k (orange) or both (blue). Spearman‘s ρ = 0.978.

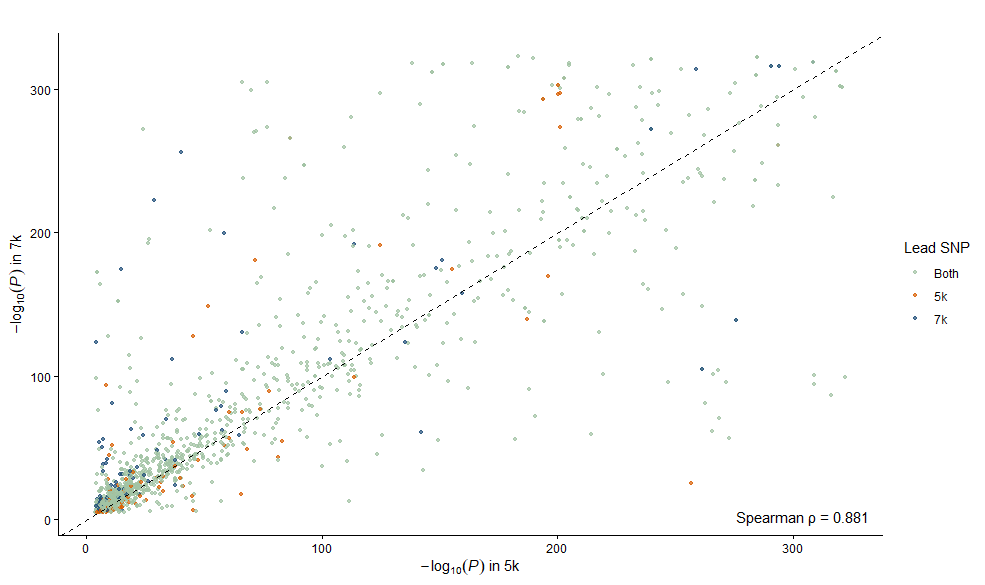

**Supplementary Figure 3.C | Correlation cis-pQTL p-values between 5k and 7k SOMAscan platforms.** Each point represents a variant-protein pair tested within the cis window and present in both datasets Points are colored according to whether the signal is detected in the 7k platform (green), 5k platform (orange) or both (blue). Spearman‘s ρ = 0.881.

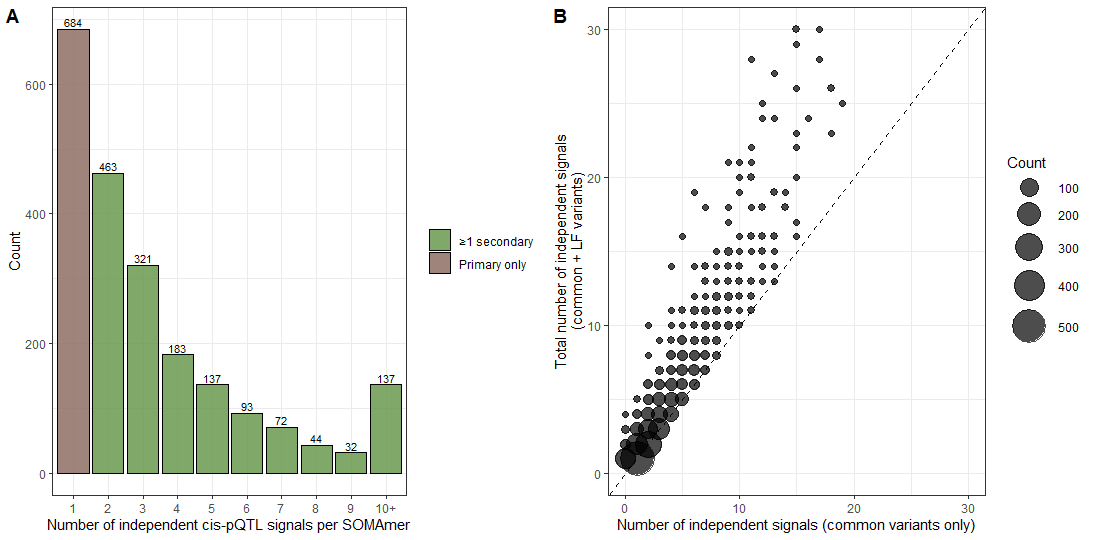

**Supplementary Figure 4 | Distribution of independent cis-pQTL signals and the contribution of low-frequency variants (A)** Distribution of the number of independent cis-pQTL signals per protein. Most proteins carry only one or a few signals, while a subset shows extensive allelic heterogeneity. Bars are coloured according to whether proteins contain only a primary signal (brown) or at least one secondary signal (green). **(B)** Comparison of the number of independent cis-pQTL signals detected using only common variants (x-axis) versus the total number detected when both common and low-frequency variants are included (y-axis). Each point represents a group of proteins with matching counts, and point size reflects the number of proteins in each group. The dashed diagonal indicates equal numbers of signals.

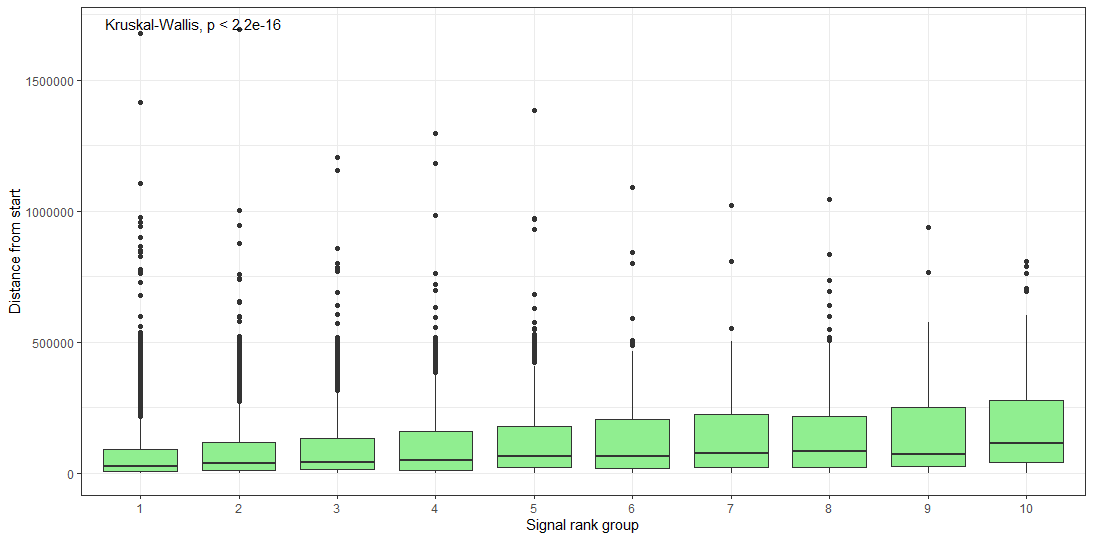

**Supplementary Figure 5 | Distance of independent cis-pQTL signals from gene boundary by signal rank.** Boxplots show the distribution of absolute distance from the gene start (bp) across signal rank groups (1 = primary signal; higher ranks indicate secondary signals). Higher-ranked signals are progressively located further from the gene start. Differences across rank groups were assessed using a Kruskal–Wallis test (P = 1.25 × 10^-100^). Boxplots indicate median value, 25th and 75th percentiles. Whiskers extend to smallest/largest value of no further than 1.5 interquartile range. Outliers are not shown.

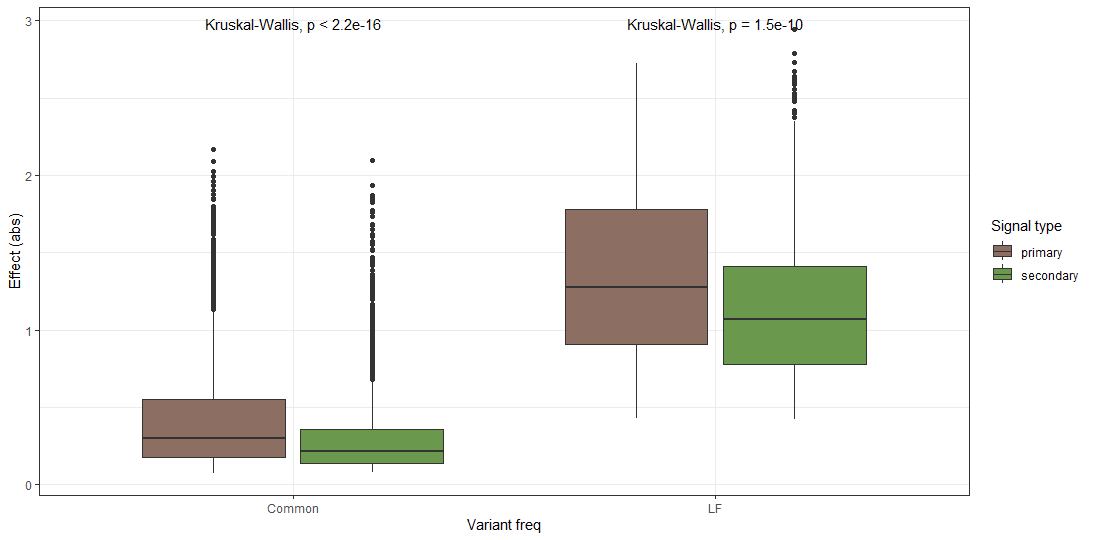

**Supplementary Figure 6 | Effect sizes of common and LF cis-pQTLs across signal type.** Absolute effect size estimates (β) stratified by variant frequency class (common vs LF) and by signal type (primary (brown) vs secondary (green)). Boxplots indicate median value, 25th and 75th percentiles. Whiskers extend to smallest/largest value of no further than 1.5 interquartile range. Outliers are not shown.

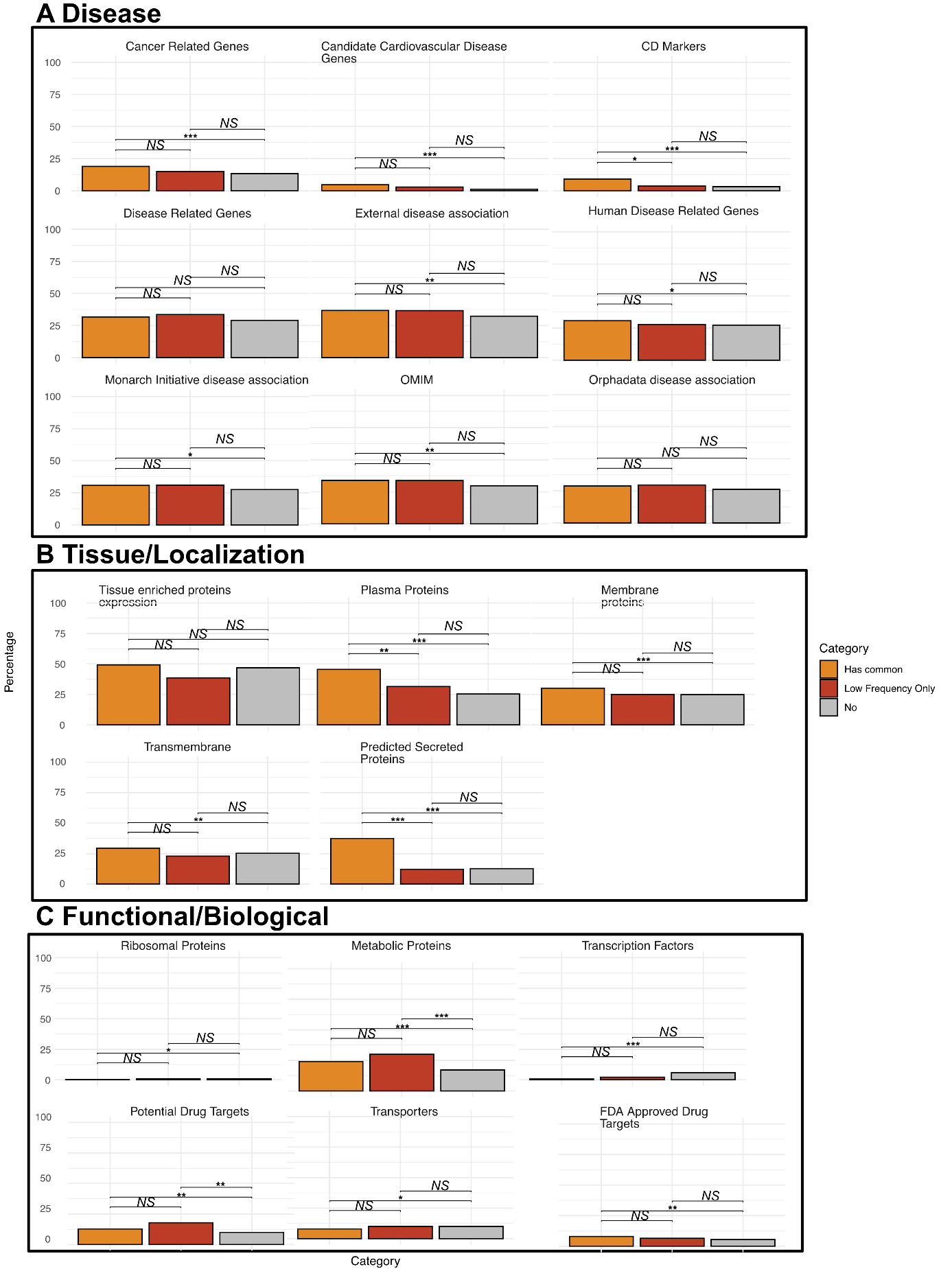

**Supplementary Figure 7 | Protein class enrichment by cis-pQTL frequency category.** Bars show the percentage of proteins in each category annotated with the indicated feature. Features are organized into three conceptual groups: (A) Disease-related annotations, including curated disease gene sets and external disease association databases (B) Tissue and localization features, including tissue-enriched expression and predicted protein localization and features and (C) Functional and biological classes, including metabolic proteins, transcription factors, ribosomal proteins, transporters, and drug target annotations. Percentages indicate the fraction of proteins within each class and statistical significance was assessed by pairwise Fisher’s exact tests, with Benjamini–Hochberg correction for multiple testing (*p < 0.05, **p < 0.01, ***p < 0.001; *NS*, not significant).
